# Patterns of litigation in France during two decades of recovery of a large carnivore

**DOI:** 10.1101/2022.10.11.511781

**Authors:** Guillaume Chapron, Gavin Marfaing, Julien Bétaille

## Abstract

The recovery of large carnivores in Europe’s human dominated landscapes is an unexpected conservation success. In France, where the wolf disappeared in 1937, the species population is now approaching one thousand individuals after the species naturally returned in the country in 1992 from Italy. Large carnivores in Europe are protected by several legal instruments, ranging from international law, to European, national or regional laws (in federal countries). There has been a limited attention allocated to how this legal protection is in practice activated in Member States of the European Union. In particular, there is little research on the role of public interest environmental litigation for large carnivore conservation. We take the example of the wolf (*Canis lupus*) in France and describe wolf-related litigation in the country during two decades. We compiled a database of case law decisions (i.e. court rulings) relating to administrative litigation about the protection of the wolf and collected a total of 275 court rulings. We found that wolf litigation occurred unsurprisingly more often in administrative courts located in regions where wolves first returned (i.e. South-East of France). Animal welfare or protection associations were the most active and successful plaintiffs. The State administration represented by its Préfets was also a plaintiff in lawsuits against illegal culling decisions made by mayors. The Préfet des Alpes Maritimes and the Minister of the Environment were regular defendants for decisions to cull wolves that were litigated by nature protection associations. Nature protection associations overall had a case winning rate higher than 50%. There were no immediately obvious inter-annual trends in wolf litigation. Our database did not allow us to quantify the total number of wolves that were effectively protected from culling decisions because court rulings made after the execution of administrative decisions did not specify whether the animals were killed or not. Bet it as it may, nature protection associations appear to conduct legally relevant litigation in view of the high success rate they achieve and conservation lawsuits belong to the portfolio of available conservation instruments.

## Introduction

The recovery of large carnivores in Europe’s human dominated landscapes is an unexpected conservation success (Chapron et al., 2014). This recovery stands in sharp contrast with global biodiversity trends that reveal an ongoing mass extinction (Ceballos et al., 2017, 2015) and repeated failures to achieve conservation targets (Xu et al., 2021). Within these trends, large carnivores are, in many places, not different from other taxa (Ripple et al., 2014). For example, lion (*Panthera leo*) populations have severely declined everywhere in Africa except in four southern countries and may disappear from many parts of the continent in the future (Bauer et al., 2015). Broadly speaking, the conservation of large carnivores is particularly challenging as these species occur at low densities, often display large home ranges and enter into conflicts with humans, their properties and activities. It is therefore remarkable that these species have recovered in a largely human-dominated and industrialized continent. Today, wolves (*Canis lupus*) occur in all continental European countries (with the exception of the Vatican, Monaco, San Marino and possibly Andorra). There are breeding populations in Denmark (Sunde et al., 2021), Belgium and the Netherlands. Italy hosts approximately 3300 wolves (Aragno et al., 2022). During the past decade wolves have made a dramatic comeback in Germany (Reinhardt et al., 2019), displaying some of the largest annual growth rates reported in wolf populations (Marescot et al., 2012). Wolves have also returned in Fennoscandia albeit in a more limited scale. There are multiple reasons why large carnivores have been able to recover in Europe. Often mentioned are a favourable public opinion generally supportive of nature conservation, an abundant prey base with high ungulate densities and a solid legal protection (Chapron et al., 2014).

Large carnivores benefit in Europe from several legal instruments, ranging from international law, to European, national or regional laws (in federal countries). Large carnivores are first listed as protected species under Article 6 of the Bern Convention on the Conservation of European Wildlife and Natural Habitats (signed in 1979, ratified by the European Union in 1982 and by France in 1990). Although some Parties to the Convention have attempted to weaken this protection, the wolf remains listed in the Annex II of strictly protected species. Enacted in 1992 as a transposition of the Bern Convention in European law, the Habitats Directive 92/43/EEC grants legal protection to all large carnivore species occurring on the EU territory of Member States. Among others, the Habitats Directive mandates that member states have protected species reaching favourable conservation status. Article 2 of the Directive states that “*measures taken pursuant to this Directive shall be aimed at ensuring the maintenance or restoration to a favourable conservation status of natural habitats and species of wild fauna and flora of Community interest*”. Species and the associated obligations and prohibitions regarding conservation and management are defined in the Directive annexes. Wolves for example are listed in most Member States in Annex II (requiring the designation of protected areas for the species) and Annex IV a) (requiring the species strict protection under Article 12 of the Directive). In some countries, or part of countries, wolves can also be listed in Annex V (instead of IV) allowing a less strict protection and the possibility of regular hunting. The strength of the Habitats Directive lies in its belonging to the legal architecture of the European Union (EU) and to the possibility of countries being sanctioned by the European Court of Justice (ECJ) which acts as a supreme court of the Union, notably through infringement procedures initiated by the European Commission. This stands in contrast with the Bern Convention where violations of that Convention are decided by agreement of the Parties, with limited enforcement of corrective measures, i.e. under Article 14(1) of the Convention, the Standing Committee, which includes all Contracting Parties, make “recommendations” to the Parties. The Habitats Directive meanwhile requires a transposition into national law. Article 188 of the Treaty on the Functioning of the European Union provides that a Directive “*shall be binding, as to the result to be achieved, upon each Member State to which it is addressed, but shall leave to the national authorities the choice of form and methods*”.

In France, the Habitats Directive has however had a minimal impact on species protection law. The Birds Directive and Bern Convention had already been implemented when the Habitats Directive came into force, and the 1976 French Act on Nature Protection guaranteed a comprehensive protection for species. Although favourable conservation status is listed as a target to be attained in Article 2 of the Habitats Directive, it is not directly included in French domestic law. Instead, Article L. 110-3 of the French Environmental Code, which appeared in the 2016 Law for the Reconquest of Biodiversity, Nature and Landscapes calls for the creation of a national biodiversity strategy, broken down into regional strategies, that “*contributes to the integration of biodiversity conservation and sustainable use objectives into public policies.*” Therefore, only the “*objectives of conservation and sustainable use of biodiversity*” are mentioned, not favourable conservation status itself. This is important because while there is a growing scholarship on the role and importance of the Habitats Directive for species conservation in the EU (Born et al., 2014), less attention has been allocated to how national protection laws are in practice activated in Member States. In particular, there is limited research on the role of public interest environmental litigation for species conservation. This article is a contribution to fill this gap. We take the example of the wolf in France and describe wolf-related litigation in the country during two decades.

### Legal conservation of wolves in France

The wolf disappeared from the French metropolitan territory in 1937 (de Beaufort, 1988). In 1992, several individuals naturally came back from Italy in Mercantour National Park and the population has since been growing (Louvrier et al., 2018). The last census reports a population of 921 wolves by the end of winter 2021/2022 (Préfet de la région Auvergne-Rhône-Alpes, 2022).

The wolf recovery has created a conflict with the local sheep farming industry which was already economically vulnerable (Chapron and López-Bao, 2014). Authorities have substantially invested into livestock protection and compensation of losses to facilitate coexistence (Gervasi et al., 2021). Since the wolf had disappeared from French territory before WWII, it was not covered by a specific legal regime until it was listed as a protected species by an executive decree in 1993 (Loubert-Davaine, 2004). That decree was annulled for formal defects in 1996 to be replaced the same year by another decree and finally by another decree in 2007 fixing the list of protected terrestrial mammals (Prieur et al., 2019). While the wolf is a protected species, this protection is not without possibilities of derogations, such as for example to cull wolves to limit damages on livestock (Audrain-Demey, 2016; Thierry, 2014).

In the Habitats Directive, Article 16 defines the conditions to grant derogations to strict protection (Epstein et al., 2019a). According to this provision, two cumulative conditions are that there must be no satisfactory alternative and that the derogation must not be detrimental to the maintenance of the population of the species in question at a favourable conservation status in their natural range. The derogation decision must then be based on one of the following grounds: protection of fauna and flora, prevention of damages (especially to livestock, concerning the wolf), public health and safety, any reasons of overriding public interest, including social and economic ones, research or education, and the taking or keeping of certain specimens on a selective basis and to a limited extent. In French law, these conditions are found in the provisions laid down by article L. 411-2, I., 4° of the Environmental Code. An executive order of 23^rd^ October 2020 sets the details of the conditions and limits under which derogations may be granted concerning the wolf. In practice, the procedure for derogating from the full protection of species is devolved with this competence belonging to the Préfets since 2007. The Préfets represent the government in a region and are the guarantors of the proper functioning of the administration and the legality of the acts of local authorities. The local services of the various ministries are also placed under the Préfet’s authority.

Decisions to derogate to species protection or decisions allegedly violating species protection can be litigated in courts. In France, the administrative court system has three layers consisting of “tribunaux administratifs” (first instance administrative courts), “cours administratives d’appel” (second instance courts of appeal) and the “Conseil d’État” (administrative supreme court). Access to justice in environmental matters is relatively broad (Bétaille, 2018; “European e-Justice Portal - Access to justice in environmental matters,” 2022). With specific reference to environmental protection associations, access to justice is governed by the French Environmental Code under Article L. 141-1 et seq. In order to obtain approval for acting as a plaintiff, article R. 141-2 provides that an environmental protection association must have been in existence for at least three years from the date of its declaration, demonstrate that it has a statutory purpose in the field of environmental protection, that it has a sufficient number of members, that it is non-profit, that it operates in accordance with the bylaws of the association, and that there are guarantees of regularity in terms of finance and accounting. The second paragraph of Article L. 142-1 provides that approved associations “*have an interest in acting against any administrative decision directly related to their purpose and their statutory activities and producing harmful effects for the environment in all or part of the territory for which they are approved, provided that this decision was taken after the date of their approval*”. Approval facilitates access to justice by presuming standing of the approved association. However, a non-approved association may still “*bring proceedings before the administrative courts*”, although it faces the obligation of demonstrating its standing in court, which is assessed in a relatively generous way by judges. In France, public interest environmental litigation has become an important part of the activities of environmental associations as well as an accepted dimension of environmental policies. It is somewhat expected by authorities that many of their decisions will be litigated and in the French administrative law tradition, litigation is part of the rule of law. Within the domain of species protection, the wolf is good illustrative example of the scope of public environmental interest litigation since the species recovery has been controversial.

### Data collection and coding

We compiled a database of case law decisions (i.e. court rulings) relating to administrative litigation about wolves. Contingently on the accessibility of these case law decisions, our goal was to build a database as exhaustive as possible. We excluded criminal litigation (e.g. wolf poaching cases), due to the lack of public availability of decisions. With permission from the Conseil d’État, we searched the intranet database of French administrative jurisdictions up to March 2019 for wolf litigation initiated by legal persons (nature protection associations, farmer organizations, different public authorities) in administrative tribunals, administrative courts of appeal and the Conseil d’État since 1997. Our database contains a total of 275 court rulings.

We coded each decision by documenting the court by which it was made, the case number and its date, the type of appeal, plaintiffs and defendants, the topic that was litigated, the number of wolves concerned by the appealed administrative decision (when documented in the ruling) and the type of decision by the court (withdrawal of the plaintiff, dismissal of the case, suspension, annulment, rejection or rejection in cassation). We also documented the ground for the decision. Finally, we recorded whether judges mentioned the following points in their rulings: availability (or lack thereof) of alternatives to culling, favourable conservation status, grounds called to derogate under Article 16(1) of the Habitats Directive and any use of scientific data in their legal reasoning.

Our analysis is descriptive only and does not aim to test hypotheses regarding the observed trends in litigations and associated wolf population dynamic, which will be the scope of separate studies.

### Patterns of litigation

For a total of N=275 cases, a quarter and the majority of them were handled by the tribunal administratif de Marseille (N=71), followed closely by the tribunal administratif de Grenoble (N=59) and then the tribunal administratif de Nice (N=49) (Table 1). The Conseil d’État was the fourth most requested court and handled 34 cases. Other courts handled significantly lower numbers of cases (less than 20).

**Table 1:**
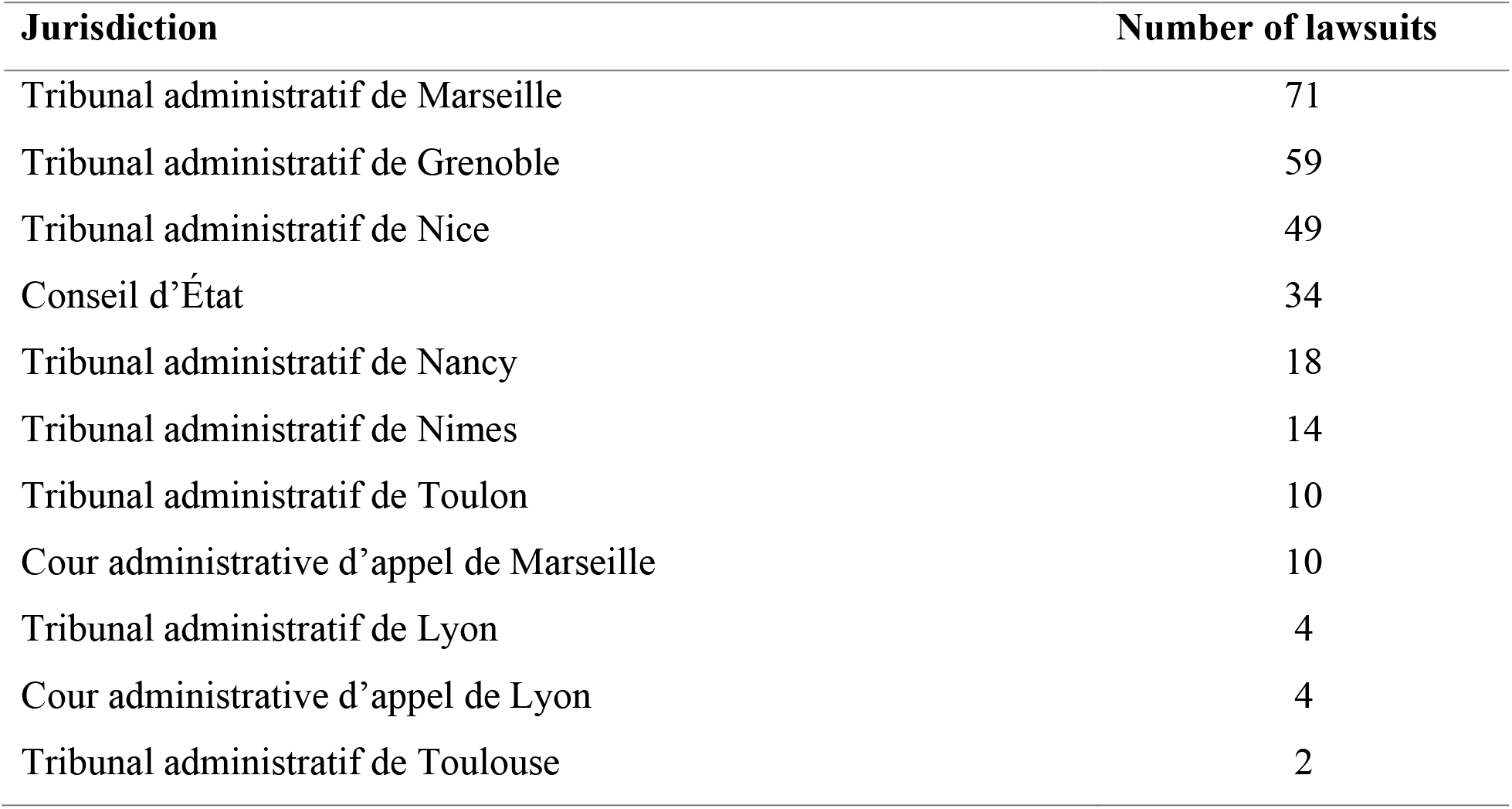
Numbers of wolf lawsuits handled by jurisdictions in France during the period 1997– 2019/03.

| **Jurisdiction** | **Number of lawsuits** |
| --- | --- |
| Tribunal administratif de Marseille | 71 |
| Tribunal administratif de Grenoble | 59 |
| Tribunal administratif de Nice | 49 |
| Conseil d'État | 34 |
| Tribunal administratif de Nancy | 18 |
| Tribunal administratif de Nîmes | 14 |
| Tribunal administratif de Toulon | 10 |
| Cour administrative d'appel de Marseille | 10 |
| Tribunal administratif de Lyon | 4 |
| Cour administrative d'appel de Lyon | 4 |
| Tribunal administratif de Toulouse | 2 |

The French Society for Animal Protection (Société Protectrice des Animaux – SPA) was the most active plaintiff, being involved into 145 cases, followed closely by the anti-hunting / nature protection Association for the Protection of Wild Animals (Association pour la Protection des Animaux Sauvages – ASPAS) (N=139) (Table 2). Conservation organizations such as Ferus (focusing strictly on large carnivore conservation), the French Federation of Environmental Associations (France Nature Environnement – FNE) and the French antenna of Birdlife International (Ligue pour la Protection des Oiseaux – LPO) were involved in less than 40 cases (Table 2).

**Table 2:** Numbers of wolf lawsuits initiated by environmental association plaintiffs during the period 1997–2019/03 (not showing plaintiffs with less than 5 lawsuits during this period). The total number of cases initiated by nature protection associations was 181, many cases had multiple association plaintiffs.

| **Plaintiff** | **Number of lawsuits** |
| --- | --- |
| Société Protectrice des Animaux (SPA) | 145 |
| Association pour la Protection des Animaux Sauvages (ASPAS) | 139 |
| One Voice | 43 |
| Ferus | 39 |
| France Nature Environnement (FNE) | 34 |
| Ligue pour la Protection des Oiseaux (LPO) | 23 |
| Groupe d'Études des Mammifères de Lorraine | 11 |

The Préfet des Hautes-Alpes was the most frequent administrative plaintiff (N=37) in court cases where the State administration litigated against decisions by mayors of communes to cull wolves (Table 3). Mayors in France do not have the devolutionary power to order wolf culling so their decisions were illegal but there needed to be a court ruling to suspend or cancel those decisions. The Préfet de la Savoie was the second most frequent plaintiff with 18 cases (Table 3).

**Table 3:** Numbers of wolf lawsuits initiated by State administration plaintiffs during the period 1997–2019/03.

| **Plaintiff** | **Number of lawsuits** |
| --- | --- |
| Préfet des Hautes-Alpes | 37 |
| Préfet de la Savoie | 18 |
| Préfet du Gard | 3 |
| Préfet du Var | 2 |
| Préfet de la Lozère | 2 |
| Préfet de l'Isère | 2 |
| Préfet des Alpes-Maritimes | 1 |
| Préfet de la Drôme | 1 |

The most frequent defendants were communes (N=70), which had made the above-mentioned decisions about wolf culling without having the authority to do so (Table 4). Closely following was the Préfet des Alpes Maritimes (N=68) and the Minister of the Environment (N=25), for decisions to cull wolves that were litigated by environmental organizations (Table 4). We also find the Prime Minister (the head of the executive power in France) was one time a defendant in an appeal by livestock farmers who demanded the Prime Minister to initiate a revision of the Bern Convention to remove the wolf from the list of protected species and for the State to take urgent measures to cull wolves (Conseil d’État, 9 May 2018, *Collectif national de préservation des activités agropastorales et rurales et autre*, n° 402013).

**Table 4:** Numbers of wolf lawsuits per defendants during the period 1997–2019/03.

| Defendant | Number of lawsuits |
| --- | --- |
| Communes | 70 |
| Préfet des Alpes-Maritimes | 68 |
| Ministre de l'environnement | 25 |
| Préfet des Hautes-Alpes | 17 |
| Préfet de la Drôme | 17 |
| Préfet des Alpes de Haute-Provence | 16 |
| Préfet de l'Isère | 12 |
| Préfet de la Lozère | 10 |
| Préfet du Var | 8 |
| Préfet de Meurthe-et-Moselle | 8 |
| Préfet des Vosges | 7 |
| Préfet de Savoie | 7 |
| Préfet de Meurthe-et-Moselle et des Vosges | 4 |
| Préfet de la Meuse | 3 |
| Préfet de l'Ardèche | 3 |
| Préfet de l'Aveyron | 2 |
| Premier ministre | 1 |
| Préfet de la région Auvergne-Rhône Alpes | 1 |

As a consequence of litigation in annulment by nature protection associations, a total of 26 wolf culling administrative decisions were annulled (i.e. cancelled) out of 48 cases. More specifically, 24 administrative decisions were cancelled on the basis of “internal legality” (i.e. the substantive rules that an administrative decision must respect in order to be lawful, e.g. breach of the law, illegality of the reasons, misuse of power) and 2 administrative decisions on the basis of “external legality” (i.e. lack of jurisdiction, formal error or procedural defect).

French law also contains a provision for fast-tracking decisions: the “référé-suspension”, with two conditions for admissibility: concerns of urgency and doubt regarding the lawfulness of the administrative decision. In the absence of urgency, the court may dismiss the case without even considering the lawfulness of the challenged decision. As a consequence of “référés-suspension” litigation by nature protection associations, a total of 38 out of 69 wolf culling administrative decisions were suspended. On the other hand, 11 complaints in “référés-suspension” were rejected on the basis of an absence of urgency and for those were concerns of urgency were not deemed absent, 20 complaints were rejected on the basis of an absence of doubt about the lawfulness of the decision (Figure 1).

**Figure 1:**
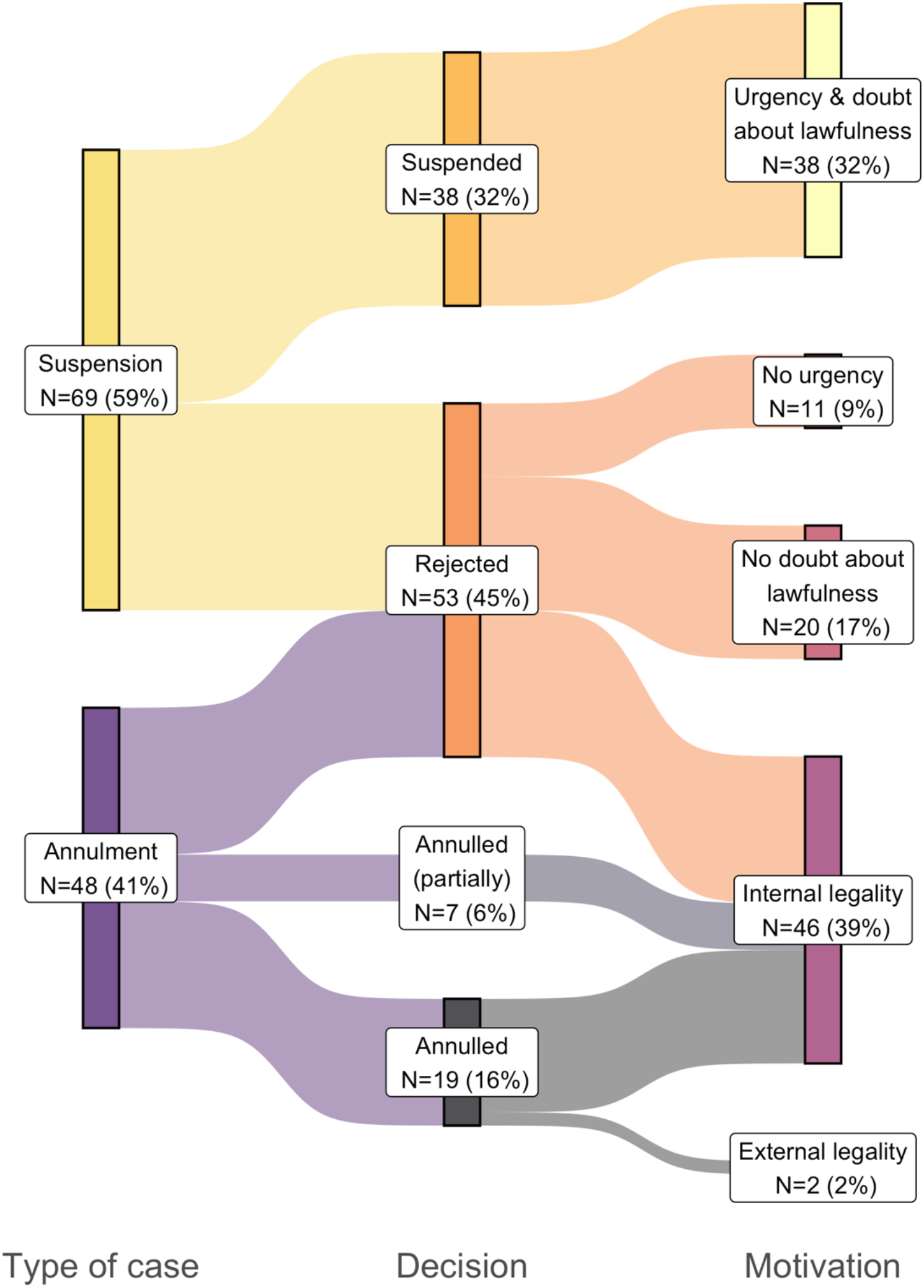
Types, decisions and motivations of wolf related court rulings during the period 1997–2019/03. The sum of decisions shown is smaller than the total number of cases initiated by nature protection associations (N=181) because some cases were withdrawn by plaintiffs or non-longer in need of a ruling (i.e. “non-lieu à statuer”).

Nature protection associations achieved an overall success rate of 55% in wolf litigation (excluding cases where they withdrew their complaints or where the courts issued a “non-lieu”, as such cases had neither a favourable nor unfavourable outcome) (Table 5). The anti-hunting / nature protection Association for the Protection of Wild Animals (ASPAS) was the most successful plaintiff (success rate of 58%), followed closely by the French Society for Animal Protection (SPA) (success rate of 57%).

**Table 5:** Success rate of nature protection association plaintiffs in wolf lawsuits during the period 1997–2019/03 (not showing plaintiffs with less than 5 lawsuits during this period).

| **Plaintiff** | **Success rate** |
| --- | --- |
| Association pour la Protection des Animaux Sauvages (ASPAS) | 0.58 |
| Société Protectrice des Animaux (SPA) | 0.57 |
| France Nature Environnement (FNE) | 0.57 |
| All associations | 0.55 |
| Ligue pour la Protection des Oiseaux (LPO) | 0.47 |
| Ferus | 0.46 |
| One Voice | 0.39 |
| Groupe d'Études des Mammifères de Lorraine (GEML) | 0.3 |

The annual number of cases was highly variable with no obvious pattern (Figure 2). Several years (2014–2016) had high number of cases (N>20) while other years (2003, 2009 or 2012) had none (Figure 2).

**Figure 2:**
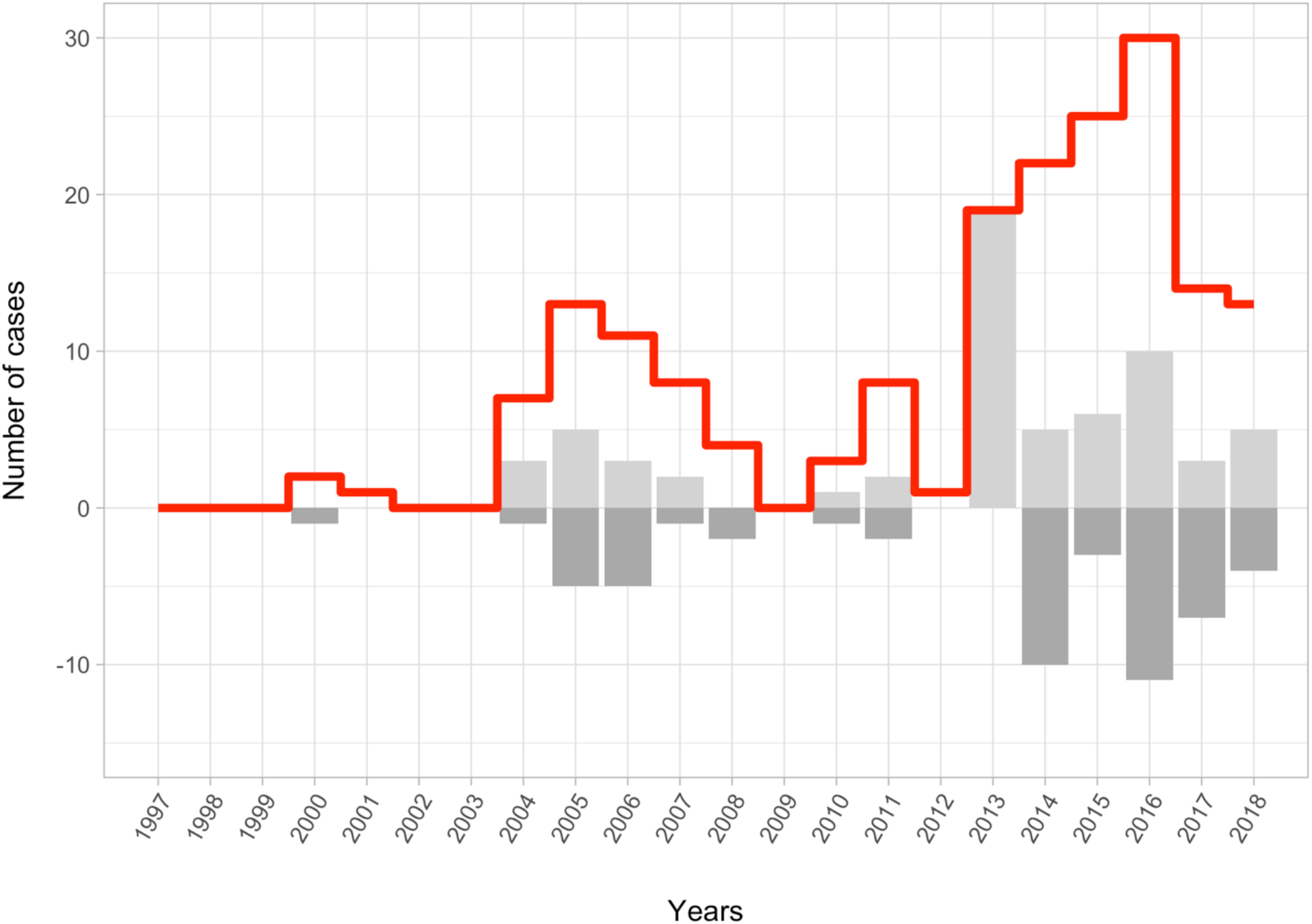
Annual number of wolf lawsuits by nature protection associations during the period 1997–2019/03. The red line shows the total number of lawsuits initiated by associations. The light grey bars indicate the annual number of court rulings where the administrative decisions sued by associations were annulled, partially annulled or suspended. The dark grey bars indicate the annual number of court rulings where the association complaints were rejected. The grey bars do not tally to the red line because the figure excludes cases where associations withdrew their complaints or where the courts issued a “non-lieu”.

Overall, the judicial decisions that validated administrative decisions to destroy wolves correspond to a total of 30 wolves. Conversely, the total number of judicial decisions overturning administrative decisions ordering the destruction of wolves corresponds to a total of 43 wolves. This corresponds to a ratio of 1.43 wolf “not killed” for 1 wolf “killed” or a 43% “return on litigation investment rate”, with the no-kill decisions becoming relatively more frequent by the end of the time series (Figure 3). Importantly, these numbers concern only decisions for which the exact number of wolves to be destroyed was mentioned, which means that a decision to cull several wolves or a whole pack without telling how many wolves were targeted does not appear in Figure 3. The number of wolves not killed may similarly be higher because some administrative decisions may have been implemented before the courts ruled.

**Figure 3:**
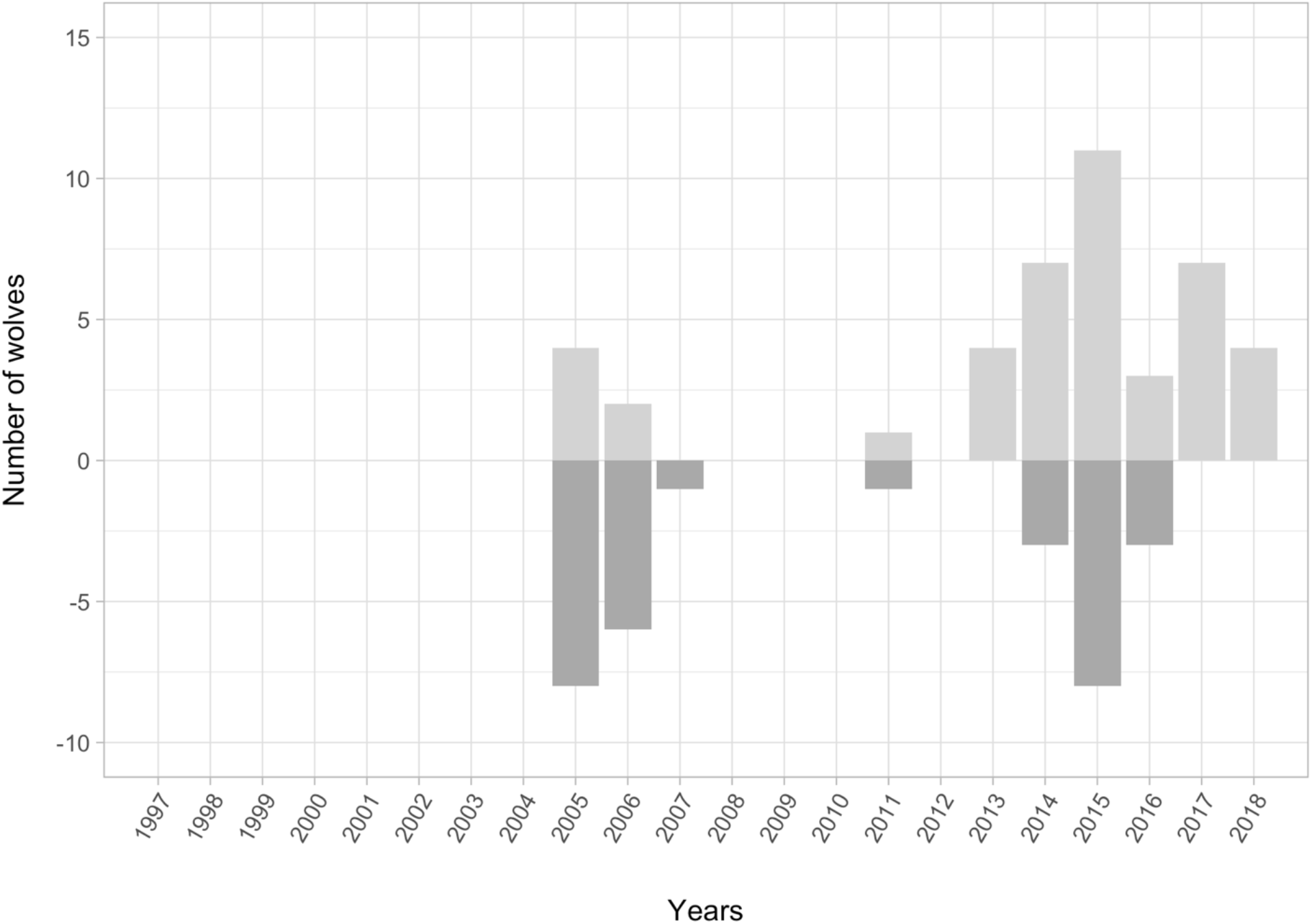
Number of wolves “ordered to be killed” (dark grey bars) or “ordered to not be killed” (light grey bars) following court cases triggered by nature protection associations during the period 1997–2019/03. This figure refers only to cases where a quota of wolves to be killed was indicated.

Regarding mentions of Habitats Directive Article 16 in court rulings, 38 decisions mentioned Article 16(1)b, of which 24 led to an annulment or suspension of the administrative decisions and 14 to a rejection.

## Discussion

Law is generally understood as being an important component of the available policy instruments to protect threatened species. It has however received a rather limited attention from conservation scholars. Writing in the scientific journal of the Society for Conservation Biology, Rohlf and Dobkin (2005) coined the term “legal ecology” and called for an increasing collaboration between the two disciplines, however their call has remained largely unanswered with their paper being cited only four times. For example, research documenting conservation responses and their impact on the status of biodiversity did not even contain the words ‘legal’ or ‘law’, as if conservation took place in an ‘alegal’ world (Hoffmann et al., 2010; Johnson et al., 2017). The available scholarship in legal conservation often focuses on the relevance of international law (Trouwborst et al., 2017a), on the interpretation of legal concepts that have a conservation relevance (Epstein, 2016; Linnell et al., 2017) or on discussing specific individual court rulings (Trouwborst et al., 2017b). In this paper we offer a new research angle to legal conservation: quantitatively describing the empirical patterns of the practice of law through public interest environmental litigation applied on the recovery of a large carnivore.

The recovery of wolves in France has, as in many other European countries, led to conflicts among stakeholders with courts having had to step in to adjudicate disagreements regarding management decisions (Epstein et al., 2019b). We found that wolf litigation occurred unsurprisingly more often in administrative courts located in regions where wolves first returned (i.e. South-East of France). The country administrative supreme court, the Conseil d’État, handled a substantial number of cases, which can be explained by the fact that it is competent in first instance over disputes concerning ministerial decisions, as opposed to prefectural decisions for which administrative courts have jurisdiction in the first instance. Cases where mayors had made decisions ordering the culling of wolves also reached the Conseil d’État after a succession of appeals.

Animal rights or welfare associations were active plaintiffs. Some of these associations (SPA) permanently employ lawyers. The most successful plaintiff, Association for the Protection of Wild Animals (ASPAS), is a sort of public-interest anti-hunting / environmental litigation association with the majority of its activities consisting of litigation. The overall high success rate enjoyed by associations was the consequence of the two most prolific plaintiffs also being the two most successful plaintiffs (58% for ASPAS with 139 appeals and 57% for SPA with 149 appeals) (Tables 2 and 5).

The State administration, represented by its Préfets, was also a plaintiff in lawsuits against culling decisions made by mayors. That the State appeared both as plaintiff and defendant might suggest that it acts in practice as a balancing force between nature conservation interests (when opposing wolf culling decided by mayors) and agricultural interests (when allowing for wolf culling). However, it is rather that these former lawsuits were initiated by the State administration to oppose a violation of the legal order since mayors did not have jurisdiction to make these decisions and not for the purpose of wolf conservation *per se*. Consequently, communes were the most frequent defendants (Table 4), being sued by both the State administration and nature protection associations.

Our analysis can also illustrate how French judges ruled on wolf lawsuits. French judges first analyse the first two conditions for derogation (the existence of alternative to destruction and maintenance in a favourable conservation status). If there are no alternatives and the decision does not harm the favourable conservation status, the judge will then examine the lawfulness of the reason for derogation. For example, the existence of alternatives (such as livestock protection) makes it unnecessary to analyse the status of the favourable conservation status and the reason for the derogation, as this is sufficient to pronounce an annulment. As a consequence, there are only 38 decisions that mention a ground for derogation, of which 24 have led to an annulment or a suspension. When citing the Habitats Directive, judges only referred to 16(1)b allowing the killing of protected species to prevent damages to e.g. livestock. There were no other Article 16 grounds used by judges which may be explained in many cases by judges only quoting French law, without explicitly referring to the Directive.

The French “Référés-suspension” is a type of order on interim measures where the litigant can obtain the suspension of an administrative decision, only if the judge considers that there is an emergency and that, at that stage of the procedure, there is a serious doubt about the lawfulness of the decision. As 11 of the 69 “référés-suspension” were rejected on the grounds of lack of urgency, it can be considered that this legal condition has had an impact on the effectiveness of this type of litigation. The relevance of this condition may thus be questioned in relation to a species protected under Annex IV of the Habitats Directive for which urgency could be presumed.

There were no immediately obvious inter-annual trends in wolf litigation. While the French wolf population has been growing almost steadily since 2000, a simple visual observation of the number of wolf lawsuits suggests there is no immediate correlation with wolf numbers. Variations in litigation may be explained by variations in the number of wolves authorized for destruction in different years (i.e. there needs to be a wolf culling decision made for a court case to happen). However, our data did not allow us to investigate this hypothesis further as we did not have access to individual wolf culling decisions (more than 1600 culling decisions were made by the Préfets just for the year 2018, the last full year of our study, for a total of 47 wolves culled (DREAL Auvergne-Rhône-Alpes, 2022)). Similarly, it was not possible to distinguish between judgments that were rendered before the administrative decisions were implemented and those that were rendered after those decisions were implemented. Therefore, our data may record a wolf as “not killed” while the animal might have been destroyed during or before the judgment proceedings. The reason is that in France, judgements made after the execution of administrative decisions do not specify whether the animals were killed or not. It is therefore impossible to make exact inferences on the demographic conservation impact of wolf litigation from this dataset. Although wolf litigation can only help wolf conservation if a given wolf is actually saved from destruction by a court ruling, it can nevertheless be said at this stage that environmental associations are conducting legally relevant litigation in view of the high success rate they achieve in courts against administrative wolf culling decisions.

Broadly speaking, one would therefore need to be careful in inferring that, when reporting a population increase after successful litigation or legal changes, the latter caused the former. Biological populations are complex systems reacting to numerous factors and often with non-linear feedbacks (Hobbs et al., 2012). This can be illustrated by another wolf population in Europe. The Swedish wolf population has been recovering since the country joined the EU in 1995, and therefore since the wolf became strictly protected in Annex II and IV of the Habitats Directive (Epstein, 2017). A simple analysis would infer that European legal protection caused this recovery. However, it was instead caused by the natural arrival of genetically unrelated animals from Finland in this highly inbred Swedish population (Liberg et al., 2005) while Sweden has in practice reneged its obligations under the Habitats Directive (Chapron, 2014; Darpö, 2016; Laikre et al., 2022). This recovery was therefore not the result of a legal factor but rather by an extra-legal factor, albeit the legal protection of wolves granted by the Habitats Directive may have prevented the re-extinction of the wolf in Sweden.

Be it as it may, public-interest environmental litigation is one factor deserving additional attention when understanding and explaining the recovery of species protection. Conservation operates within the laws of the country in which it happens, which for EU Member States include the Habitats and Birds Directives and conservation lawsuits belong to the portfolio of available conservation instruments.

